# Tap Dancing Frogs: Posterior Toe Tapping and Feeding in *Dendrobates tinctorius*

**DOI:** 10.1101/2023.09.15.558032

**Authors:** Thomas Q Parrish, Eva K Fischer

## Abstract

Animals have myriad adaptations to help them hunt and feed in the most efficient and effective manner. One mysterious behavior related to hunting and feeding is the posterior toe tapping behavior of some frogs. Biologists and hobbyists alike have long noticed this behavior, but there is little empirical data to explain its causes and consequences. To test the hypothesis that tapping is related to feeding and modulated by environmental context, we conducted a series of related experiments in the Dyeing poison frog, *Dendrobates tinctorius*. We first confirmed that tap rate was higher during feeding as has been observed in other species. Interestingly, this effect was heightened in the presence of a conspecific. We next asked whether frogs tapped less under conditions when prey were visible, but inaccessible. Finally, we asked whether *D. tinctorius* adjusted tap rate based on substrate characteristics and whether prey capture success was higher when tapping. In addition to confirming an association between tapping and feeding, our work demonstrates modulation of toe tapping based on social context, prey accessibility, and substrate characteristics. Based on our findings, we suggest that tapping could act to induce prey movement and thereby facilitate prey detection and capture by frogs.

**Significance Statement:** The toe tapping of some amphibians is an intriguing behavior that has attracted attention from researchers and hobbyists, yet the functional role of toe tapping remains poorly understood. Previous studies have noted an association between toe tapping and feeding in other species, and our work does so quantitatively and experimentally in Dyeing poison frogs, *Dendrobates tinctorius*. We demonstrate that frogs modulate tap rate based on the presence of conspecifics, prey accessibility, and substrate type. We speculate that – alongside other potential functions – toe tapping may act as a vibrational stimulus to facilitate prey detection and capture.

## Introduction

The toe tapping of various species of amphibians is an intriguing behavior that has attracted attention from researchers and pet owners. Variably referred to as toe tapping, twitching, wiggling, and trembling, this curious behavior is the subject of YouTube videos, TikToks, and Twitter threads. Indeed, a recent publication took advantage of online video sharing platforms to compile a toe tapping dataset of 222 videos of 389 individuals representing 63 frog species (Claessens, Ganchev, Kukk, Schutte, & Sloggett, 2020). Yet despite being readily observed and widely documented, the functional role of toe tapping remains poorly understood.

Among amphibians, toe tapping falls under the general umbrella of pedal signaling and luring, which ranges from ostentatious movements of the entire limb to subtle tapping of fingers or toes (reviewed in (Erdmann, 2017; Hödl & Amézquita, 2001)). Under this umbrella, a related set of similar-looking toe movements have been broadly discussed in the context of visual and vibrational communication. Toe tapping has been suggested to function in prey capture (Claessens et al., 2020; Hagman & Shine, 2008; McFadden, Harlow, Kozlowski, & Purcell, 2010; Sloggett & Zeilstra, 2008), mate attraction and courtship (Landestoy & Ortíz, 2015; Starnberger, Maier, Hödl, & Preininger, 2018), aggression (Bee, Reichert, & Tumulty, 2016; Furtado, Márquez, & Hartz, 2017), and has also been suggested to be a ‘displacement behavior’ with no functional relevance (Furtado et al., 2017; Furtado & Nomura, 2014).

We regularly observe Dyeing poison frogs (*Dendrobates tinctorius*) in our lab tapping their posterior toes (Movie S1 & S2). Toe tapping has been documented in various species of poison frog (family Dendrobatidae), with some bias toward observations in this group likely resulting from the fact that these brightly colored, diurnal frogs are popular as pets and in zoos and aquaria (Claessens et al., 2020). Two recent studies demonstrated increased toe tapping associated with feeding in poison frog species closely related to *D. tinctorius*. Schulte and König (Schulte & König, 2023) reported more tapping in response to both small (fruit flies) and large (crickets) prey items, but not conspecific calls, in *Dendrobates auratus*; and Vergara-Herrera and colleagues (Vergara-Herrera, Cocroft, & Rueda-Solano, 2023) report detailed tap characterization in *Dendrobates truncatus*, including accelerated tapping right before prey capture.

Poison frogs have dietary specializations for small prey items (Santos, Coloma, & Cannatella, 2003), and in the wild *D. tinctorius* feed on a variety of arthropod prey (Rojas & Pašukonis, 2019). Their prey is small and quick, and prey movement is key for detection and capture. The essential role of prey movement for prey capture in amphibians was demonstrated by studies showing that toads readily strike at any object of the appropriate size and moving in the correct direction (i.e., ‘worm like’ lines moving horizontally) (Ewert, 1987). These studies garnered frogs and toads the reputation of being stereotyped in their hunting strategies, with sensory constraints limiting their ability to plastically adjust behavior. However, recent years have seen a growing number of studies cataloging the ability of these animals to integrate complex, multi-modal sensory information (e.g. (Amézquita & Hödl, 2004; de Lourenço, Haddad, & de Sá, 2020; Deban, O’Reilly, & Nishikawaa, 2001; Hartmann, Giasson, Hartmann, & Haddad, 2005; James et al., 2022; Narins, Grabul, Soma, Gaucher, & Hödl, 2005; Toledo, Araújo, Guimarães, Lingnau, & Haddad, 2007)).

Though primarily explored in the context of predator evasion and conspecific communication (e.g. James et al., 2022; Narins, 1990; Warkentin, 2005), the exquisite vibrational sensitivity of anurans may also play a role in prey capture. Using a mechanical model, Hagman and Shine (Hagman & Shine, 2008) demonstrated that cane toads vibrate their toes close to the frequency most effective at attracting conspecific prey. Vibrational communication and seismic stimuli through surfaces have also been shown to be important for amphibian’s arthropod prey. For example, flies can feel vibrations through their legs and react to them (Jarman, 2002). Taken together, this led us to predict that toe tapping could – among other functions – be a vibrational and/or visual stimulus associated with prey capture and feeding.

Under the assumption that toe tapping indeed has a functional role in feeding and foraging, we predicted that *D. tinctorius* would adjust their tapping behavior based on environmental context. We performed a series of experiments to test this idea. First, we confirmed that *D. tinctorius* do indeed tap more while feeding. Second, we asked whether frogs tapped less under conditions in which they could see, but not capture, prey. Third, we tested whether *D. tinctorius* modulate tap rate based on substrate characteristics and asked whether prey capture success was higher when tapping. Alongside recent studies in related species, our findings contribute to our understanding of this visually conspicuous yet functionally mysterious behavior.

## Methods

### (a)Animals

*D. tinctorius* were housed in our animal facility at the University of Illinois Urbana-Champaign. Frogs were housed breeding pairs in 45x45x45 cm terraria containing soil, moss, leaf litter, live plants, coconut shelters, and a small pond. We recorded only a single frog per terraria (i.e., per pair) in any given trial. Terraria were maintained at 70-100% humidity and 22-24ºC on a 12L:12D light cycle. We fed frogs *Drosophila hydei* or Drosophila *melanogaster* dusted with vitamins three times weekly. All animal care and experimental procedures were approved by the University of Illinois Animal Care and Use Committee (IACUC protocol #20147). We used the STRANGE framework in our experimental design and note that – while toe tapping is observed in the wild – these are captive bred and reared animals where there is potential for lab selection.

### (b)Experiment 1: Tapping in feeding versus non-feeding states

In the first experiment, we tested if *D. tinctorius* tap more while feeding versus not feeding. For feeding trials, we dropped ½ teaspoon of *Drosophila* into the frogs’ terrarium. Once frogs began hunting, we took one minute of high-speed video (240 frames per second). For non-feeding trials, we took one minute of high-speed video while frogs were not feeding. All trials took place between 11:00 and 14:00. We also recorded partner proximity (frogs are housed in breeding pairs) during filming. Videos were analyzed as described below. N=12 feeding and N=10 non-feeding trials with N=8 total individuals (some frogs were tested twice).

### (c)Experiment 2: Tapping when prey is on a different surface

In the second experiment we tested whether tapping behavior was affected by prey being on a different surface and inaccessible to the frogs. We placed *Drosophila* inside a small, clear petri dish so that the frogs could see the prey but not feed on it, and the prey was not on the same surface as the frog. Petri dishes were placed in frogs home terraria during normal feeding time, and we recorded high-speed video for one minute once frogs began striking at prey. Paired trials with freely moving flies were conducted as described above the next day. We performed N=16 inaccessible prey and N=14 free-moving prey trials with N=16 individuals (two frogs were excluded on the second day because they were transporting tadpoles).

### (d)Experiment 3: Tapping variation between surfaces

In the third experiment, we asked whether tap rate depended on substrate type. We set up experimental terraria with four substrate types: soil (a natural, unpliable surface), leaf litter (a natural, pliable surface), gel (1% agar; an unnatural, pliable surface), and glass (an unnatural, unpliable surface). We chose these surfaces to obtain a variety of pliability and familiarity. We placed frogs singly into test terraria and gave them one-hour acclimation to the novel environment. Following acclimation, we fed frogs and collected data as in Experiment 1. In addition, we recorded the number of strikes frogs made in attempts to catch prey, and how many of these strikes were successful. We performed all trials one day after last feeding to control for satiety status. Each individual performed one trial on every surface, apart from two individuals performing two on each surface. N=13 trials per surface with N=11 individuals. All trials were analyzed for tap rate. For strike and success rate, three trials were excluded from analysis due to video loss.

### (e)Data Analysis

We quantified tap number from slow motion video and standardized data for each frog. To calculate the number of taps per minute, we divided the number of taps on each foot by the number of seconds the foot was visible to obtain taps per second. We then multiplied that by 60 to calculate taps per minute for each foot and added the two feet together to get total taps per minute. For experiment three, we calculated success rate by dividing the number of successful strikes by the total number of strikes. To test for potential biases introduced by toe visibility across contexts, we compared visibility rates across surfaces and found no differences (main effect of surface: F_3,48_ = 0.45, p = 0.7168).

We used R v4.4.2 (Team, 2022) in Rstudio v 3.1.446 (Posit Team, 2023) for all statistical analyses. As in previous studies (Vergara-Herrera et al., 2023), we did not find an effect of sex on tap rate and therefore omitted this variable from further analyses. In the first experiment, we tested for differences in tap rate using a linear mixed model that included feeding status and partner presence as fixed effects and frog ID as a random effect. In the second experiment, we used the same approach (excepting the random effect of individual) to compare frogs feeding on free flies (data from Experiment 1) to frogs feeding on flies confined to a petri dish. In the third experiment, we included substrate type as a fixed effect and frog ID as a random effect. For all linear models, we evaluated significance via ANOVA followed by Tukey corrected post hoc testing with the package emmeans in R. We tested for correlations between tap rate, strike rate, and strike success using Spearman correlations. We additionally tested for a substrate-dependent relationship between prey capture success and tapping, where capture success rate was predicted by surface type, tap rate, and their interaction.

## Results

### (a)Frogs tap more when feeding, especially in the presence of a conspecific

Frogs tapped significantly more when feeding versus non-feeding (main effect of feeding: F_1,17_ = 116.65, p <0.0001) and this effect was more pronounced in the proximity of their partner frog (feeding*partner interaction: F_1,18_ = 7.36, p = 0.0143) (Fig. 1a). While partner proximity increased tap rate in the presence of prey (feeding alone vs feeding with partner: t_17.8_ = -4.121, p = 0.0033), tap rate was not increased by partner proximity in the absence of prey (not feeding alone vs not feeding with partner: t_17.8_ = -0.277, p = 0.9923). The average tap rate was 389 taps/min (range: 305 - 468) when feeding alone, 684 taps/min (range: 492 - 997) when feeding near a partner, 50 taps/min (range: 4 - 149) when not feeding away from a partner, and 43 taps/min (range: 5 - 92) when not feeding but near a partner.

**Figure 1:**
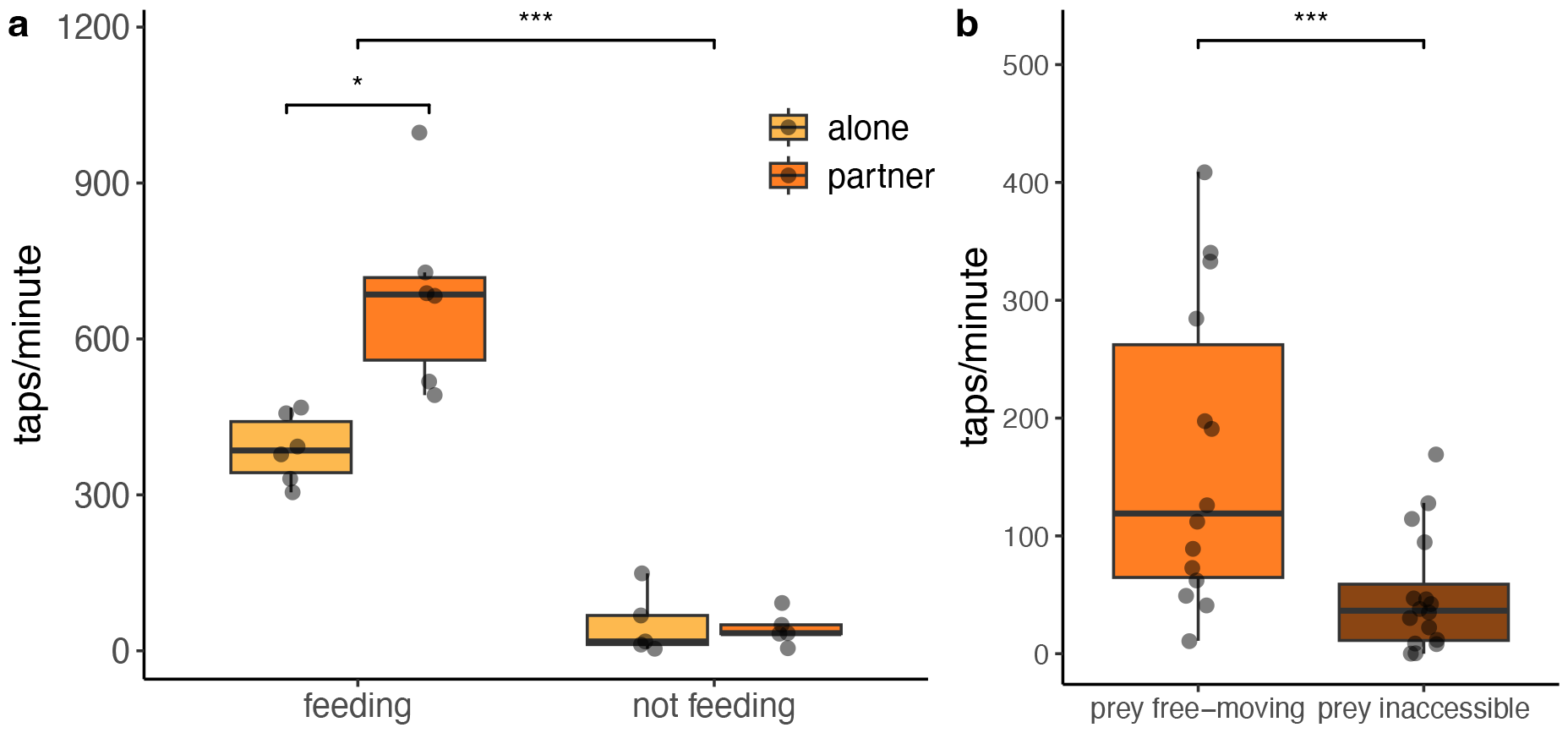
Increased tapping when feeding on accessible prey. **(a)** Frogs tapped significantly more when feeding versus not feeding, and this effect was more pronounced in the presence of a partner frog (dark orange) than when alone (light orange). **(b)** Tap rate was significantly lower when prey was on a different surface and inaccessible to frogs (flies inside a petri dish; Movie S3).

### (b)Frogs tap more when prey is accessible

Tap rate was significantly lower when prey was inaccessible and on a different surface from the frogs (F_1,14_ = 19.85, p = 0.0005) (Fig. 1b). The average tap rate when frogs could see, but not access, flies was 50 taps/min (range: 0 - 169); significantly lower than the 166 taps/min (range: 11 - 409) while feeding on free moving flies (Fig. 1b). Importantly, we observed this lower tap rate even though frogs continuously attempted to catch inaccessible prey (Movie S3).

### (c)Tap rate depends on surface type

There was a significant difference in tap rate between surface types (F_1,40.6_ = 6.21, p = 0.0014) (Fig. 2a). Average tap rates were 98 taps/min for soil (range: 0 - 417), 255 taps/min for leaf (range: 24 - 528), 118 taps/min for gel (range: 0 - 440), and 64 taps/min for glass (range: 0 - 211). Post hoc comparisons revealed that frogs tapped more on leaves than all other surfaces (gel: t_38.4_ = 2.87, p = 0.0317; glass: t_38.4_ = 4.01, p = 0.0015; soil: t_38.4_ = 3.30, p = 0.0107) (Fig. 2a). Tap rates in experimental tanks (Experiment 3) were on average lower than when frogs were in their home terraria (Experiment 1), but still higher than when not feeding in home terraria.

**Figure 2.**
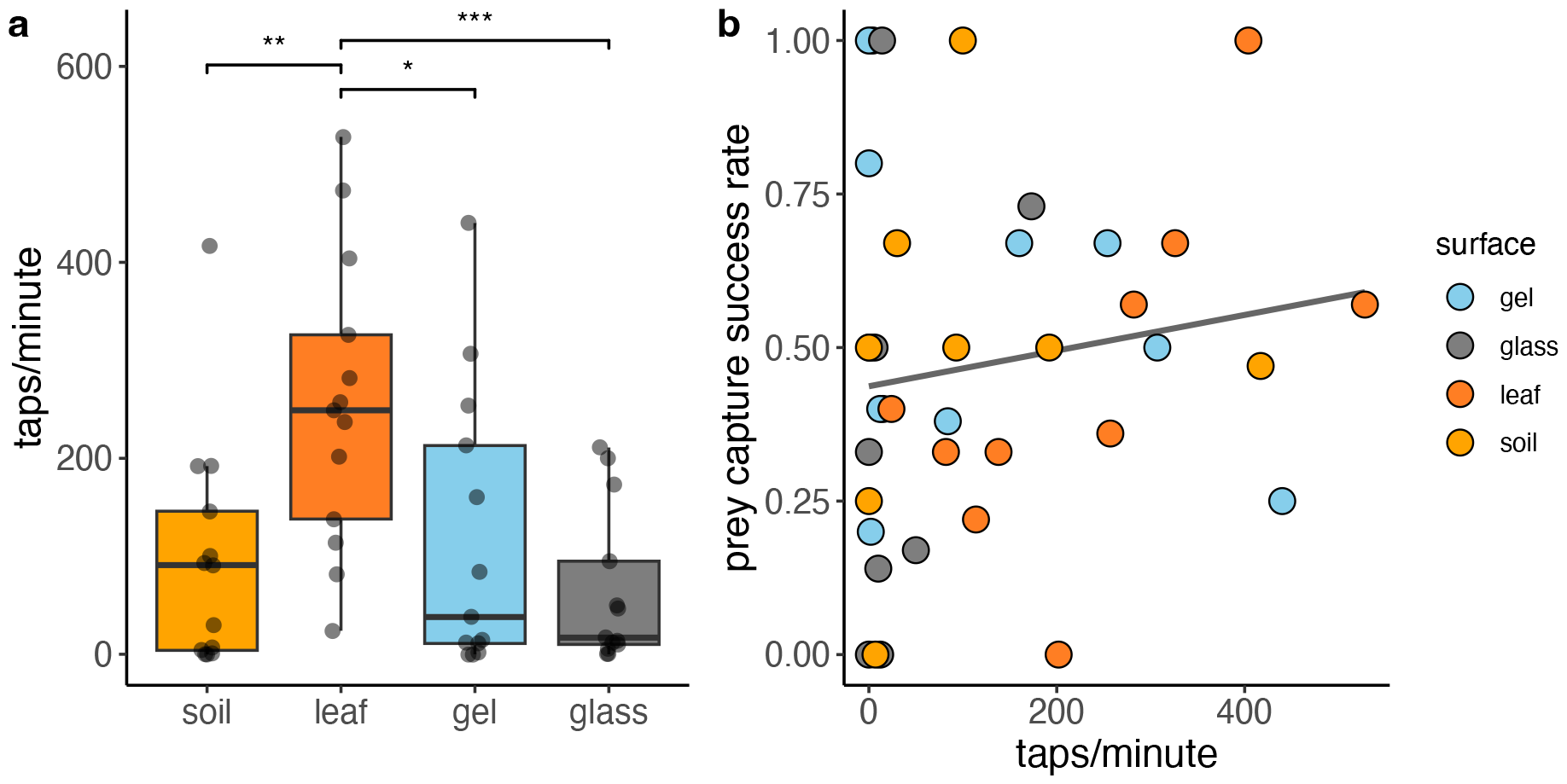
Surface type affects tap rate but not prey capture success. (a) Tap rates differed based on surface. Tap rate was higher on leaves (orange) as compared to soil (yellow), gel (blue), or glass (grey). (b) Prey capture success rate did not differ based on surface type and was not predicted by tap rate.

While tap rate differed between surface types, there was no difference in the total number of strikes (F_3,28_ = 1.41, p=0.259) or success (F_3,28_ = 0.81, p = 0.497) based on substrate. Strike rate increased with tap rate (r=0.318, p=0.045); however, tap rate did not significantly predict prey capture success overall (tap rate main effect: F_1,30_ = 2.29, p = 0.142) or based on surface type (tap rate*surface interaction: F_3,28_ = 1.43, p = 0.254) (Fig. 2b).

## Discussion

Taken together, our findings demonstrate that tapping is associated with feeding and tap rate is modulated by partner proximity, prey accessibility, and substrate characteristics in *D. tinctorius*. In line with our predictions, we found that the frogs tapped more while feeding, when prey was accessible, and on a vibrationally responsive surface. Unexpectedly, we also observed an increase in feeding-related tapping in the presence of a breeding partner. Counter to our expectations, we did not find a relationship between tap rate and prey capture success. We discuss interpretations and implications of our findings below.

An association between feeding and posterior toe tapping has long been noted by researchers and hobbyists, and recent empirical work substantiated this relationship in two closely related species of poison frog (Schulte & König, 2023; Vergara-Herrera et al., 2023). Experiment 1 confirmed that tapping is associated with feeding in *D. tinctorius*, with frogs increasing their tap rate up to 10-fold in the presence of prey. Interestingly, tap rate was further increased when feeding in proximity of a breeding partner, while partner proximity alone did not increase tap rate. At present, we do not know the reason for this social modulation of feeding-related toe tapping behavior (see further discussion below). We highlight that – at a maximum of almost 500 taps per toe per minute – this behavior is incredibly fast for any vertebrate muscular movement (Rome, 2006). In brief, these findings demonstrate an association between tapping and feeding and provide interesting avenues for further study.

Given the vibrational sensitivity of both frogs and their arthropod prey, and the importance of prey movement for prey detection, we hypothesized that toe tapping could function as a vibrational stimulus that facilitates prey capture, as has been demonstrated for other forms of toe tapping and pedal luring (Erdmann, 2017; Hagman & Shine, 2008; Sloggett & Zeilstra, 2008). To test this idea, we examined the relationship between tapping and surface type. When frogs could see but not capture flies, frogs tapped significantly less but *still hunted*. We suggest that this change in tap rate may be related to changes in vibrational stimuli and/or feedback from prey capture success. This observation suggests that frogs might alter their tapping behavior based on prey’s responses.

To test the potential connection to vibrational stimuli further, we next asked whether *D. tinctorius* modulated tapping based on surface type. We found that frogs tapped at a significantly higher rate on the leaf surface. This aligned with our prediction that tapping would be enhanced on vibrationally responsive surfaces. We were concerned that tapping behavior might be impacted by the naturalness and familiarity of each surface, and the novel environment of the test terraria. Tap rate was overall lower in the experimental terraria than in home terraria; however, frogs hunted on all test surfaces, with no differences in prey capture attempts or prey capture success. Additionally, we found that frogs tapped less on soil than leaves, despite both surfaces being familiar to frogs. Taken together, these observations demonstrate that frogs modulate tapping behavior based on substrate type and independent of prey capture attempts.

We expected higher tap rates to be associated with increased prey capture success; however, this prediction was not supported, in general or based on surface type. Similarly, we found no difference in prey capture success across surface types. We suspect this lack of a relationship may be an artifact of our feeding method in the lab, where frogs are provided a large number of flies and prey capture success is artificially high. While this possibility requires further study, our demonstration of an association between tapping behavior, capture attempts, and surface type is suggestive and opens the door for further experiments.

Additional work is also needed to clarify additional, non-mutually exclusive roles and reasons for toe tapping. One alternative explanation for tapping is a generalized displacement and/or excitatory response (Furtado et al., 2017; Furtado & Nomura, 2014). Under this assumption, tapping has no functional role but is a reflex or arousal response to food or movement, akin to a dog wagging its tail. This alternative could contribute to a lower tap rate when prey was inaccessible, assuming arousal is lower when capture is unsuccessful. However, such an effect cannot explain all our findings here, given that frogs also modulated tap rate based on surface type and independent of prey presence and movement or feeding success.

Another non-mutually exclusive alternative is that tapping may be involved in social communication between frogs, in agonistic and/or affiliative contexts (Furtado et al., 2017; Landestoy & Ortíz, 2015; Starnberger et al., 2018). In poison frogs specifically, toe tapping has been reported during feeding (Schulte & König, 2023) and observed during courtship (Barquero & Arguedas, 2022) in *D. auratus*; however, presentation of advertisement calls did not increase toe tapping in this species (Schulte & König, 2023). Tapping has similarly been noted during the later stages of courtship in *D. tinctorius* (Rojas & Pašukonis, 2019) where it may also play a role in vibrational signaling. We observed an increase in toe tapping while feeding in the presence of another frog, but this social modulation occurred only when prey was present. These observations suggest that any social function – be it cooperative or antagonistic – is in addition to, not instead of, a feeding-related role. We emphasize that our study was not designed to explicitly test a role for toe tapping in social communication and we did not monitor tapping during courtship or aggressive interactions. We note that conspecifics in our study were the breeding partners of focal individuals, and that familiarity influences aggressive and affiliative behaviors. In sum, while other non-mutually exclusive roles for posterior toe tapping exist and are of interest for future study, our data here support a connection between toe tapping and feeding that is socially and environmentally modulated in *D. tinctorius*.

### Conclusions

Taken together, our findings confirm an association between toe tapping and feeding and demonstrate that *D. tinctorius* modulate tapping behavior based on social and environmental context. The observation that frogs alter their tapping frequency – but not their strike rate – based on substrate type, suggests that tapping has a functional, context dependent role. In line with previous studies, we speculate that tapping may stir small arthropods via substrate vibrations and thereby help *D. tinctorius* detect prey. By quantifying tap number, we also confirm the impressive speed with which frogs perform the behavior. Future studies characterizing tapping biomechanics, measuring substrate vibrations caused by tapping, and testing the vibrational sensitivity of both frogs and prey will further our understanding of tap-dancing toes.

## Acknowledgements

We would like to thank Samta Oza for help with data collection, and members of the Fischer Lab for their support and feedback on previous version of the manuscript. Thanks to Phil Anderson for advice on project design and assistance with high-speed filming.

## Data Availability

Data analyzed in the current study are provided in the Supplemental Materials.

## Author Contributions

Conceptualization, data curation, analysis, investigation, visualization and writing by TQP and EKF. Methodology by TQP. Project administration, funding acquisition and supervision by EKF.

